# Surrogate Verifiers Shape Decisions in Closed-Loop Experimental Design

**DOI:** 10.64898/2026.11.18.767545

**Authors:** Charlotte Merzbacher, Jannus Goecke, Jake Cohen, Martin Patz

## Abstract

High throughput, automated screening has revolutionized biological experimentation and prompted predictions of AI-enabled “self-driving” labs. Deployed autonomous lab systems typically use a learned surrogate to predict new experimental candidates and rank which conditions deserve expensive follow-up. Determining whether a computational surrogate verifier can substitute for an experimental assay’s ground truth requires an understanding of model accuracy, uncertainty, and extrapolation. In this work, we present surrogate verifiers trained on N=7,688 *Lacticaseibacillus* growth curves exploring a 10-dimensional media component experimental design space. We train surrogate models across six distinct families including polynomial, Gaussian-process, tree-ensemble, neural, and Monod-kinetic forms to predict maximum optical density (OD600). Surrogates ranked by the most commonly reported accuracy metrics (held-out R^2^; MAE; RMSE) disagree sharply on which experiments to test next and the best achievable yield. We evaluate models in a Bayesian optimization loop and find that the model used as the active learning oracle matters more than the acquisition function or batch size. We apply conformal calibration to build a model-agnostic assessment criterion of uncertainty and model coverage. We end by evaluating model accuracy and calibration under transfer to an unseen related bacterial strain, and confirm on a prospective validation plate that the transferred verifier’s intervals fail in the direction and region our retrospective analysis predicts. This work explores the surrogate verifier within the loop as a previously underexplored axis of closed-loop design and points towards best calibration practices.

## 1 Introduction

Automated robotic labs (see Figure 1) that can perform many hundreds of experiments in parallel, paired with AI models that can suggest and evaluate candidate experiments, promise to greatly accelerate biology research [Cheng et al., 2026, Helleckes et al., 2026]. “Lab-in-the-loop” systems train a model on experimental data, then iteratively design further experiments based on that model’s ranking of possible experiments [Kitano et al., 2023]. Only the highest-ranking members of the candidate pool receive additional ground-truth verification via wet-lab validation studies; the vast majority are evaluated entirely *in silico*. In addition to ranking experiments, the ML model stands in for the ground truth wet-lab results when benchmarking competing design strategies against each other. In both roles, the model acts as a surrogate verifier, but in previous literature its reliability is assumed rather than measured [Siska et al., 2026].

**Figure 1:**
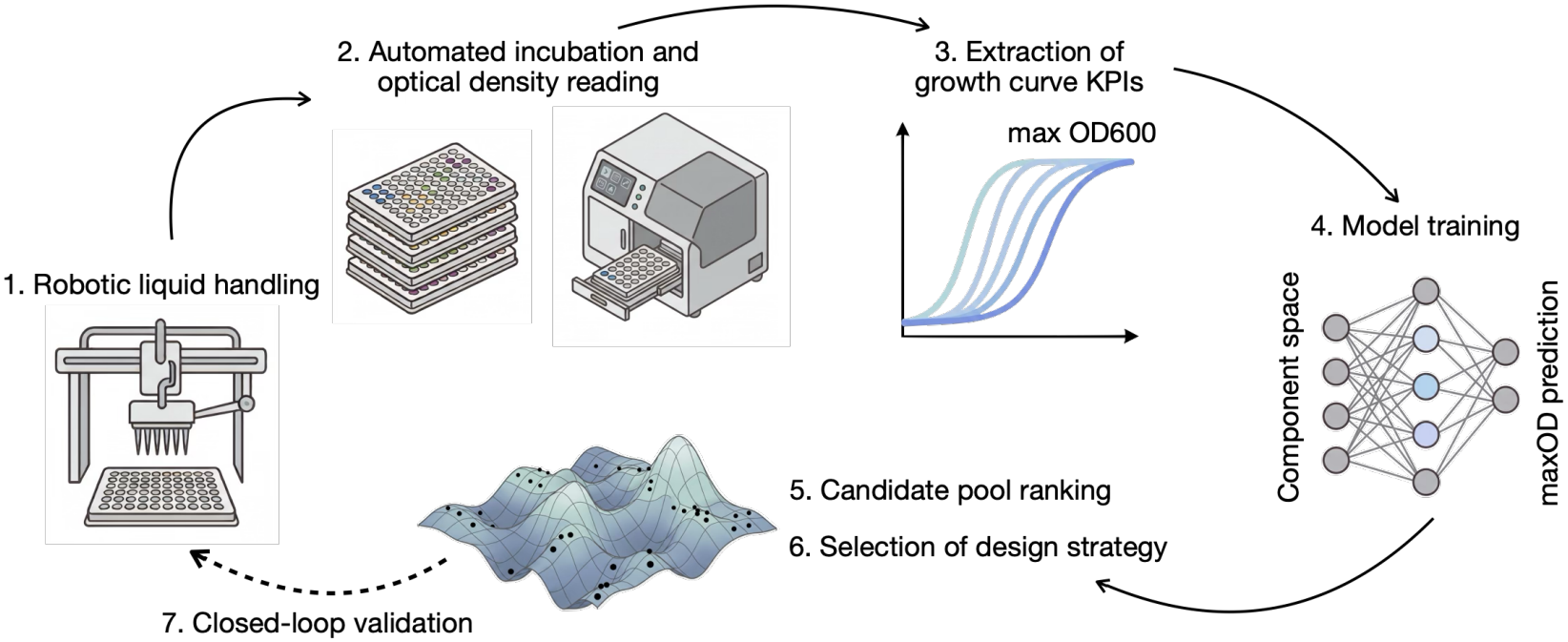
Schematic of autonomous lab. A liquid handler pipets 96-well plates with different medium conditions. These plates are then incubated for 48hrs at 37ºC with automated optical density (OD600) measurements taken via a plate reader. The maximum OD600 is extracted from the resulting growth curves and used as a regression label for supervised surrogate model training. Models are then used to assess the larger candidate pool and select the design strategy for future (dashed line) closed-loop experimental iterations.

A robust literature on ML-guided experimental design includes Bayesian optimization methods that show improved titer and sample-efficiency compared to standard design of experiment (DoE) [Lapierre et al., 2025, Cosenza et al., 2022, Narayanan et al., 2025]. The models in these active learning loops are usually selected and evaluated based on accuracy metrics, such as *R*^2^, mean absolute error (MAE), and root mean squared error (RMSE) on a held-out test set. These accuracy metrics evaluate average error over data already collected, but optimizing for high-performing conditions selects outliers by definition. Fitness for either ranking or benchmarking depends on properties such as how trustworthy the model’s uncertainty is in the region the design loop is querying, and how well beyond the observed data it extrapolates.

In this work, we examine the performance and limitations of surrogate verifiers for bioprocess media formulation optimization. We built verifier models across six model families on a dataset of N=7,688 growth curves across three related *Lacticaseibacillus* strains sharing a common 10-component media design space (5 levels per component; >9.7M experimental candidates). We aim to assess when and how surrogate verifiers can replace wet-lab experimental ground truth by investigating whether surrogates fitted on the same data agree on their rankings and predictions. We show that the growth prediction depends on the model that makes it: families closely ranked by accuracy disagree about which conditions to run next, how much growth is achievable, and about which design strategy a benchmark should recommend; we trace that disagreement to the geometry of the fitted surfaces. We use conformal calibration as a model-agnostic criterion to assess validity and certification sharpness across families. Finally, we perform transfer learning to a new strain to assess whether confidence and calibration survive transfer. We find that fine-tuning a verifier onto a new strain improves its accuracy but leaves its uncertainty miscalibrated, and assess how many experiments from a new organism are needed to guarantee calibration coverage. Taken together, these results indicate that the surrogate inside a design loop is not a neutral measuring instrument but a component of the experiment, whose choice must be reported and qualified alongside the design strategy it is used to score.

## 2 Background

### 2.1 Active learning for bioprocess engineering

Machine learning-guided experimental design is an established practice in bioprocess development [Karimi Alavijeh et al., 2024]. Bayesian optimization is the most widely applied method and has been used in optimizing microbial growth media [Lapierre et al., 2025], cell-culture media [Narayanan et al., 2025, Cosenza et al., 2022], chromatography [Pham et al., 2025], and cell therapy manufacturing [Claes et al., 2024], reporting comparable or improved titer at a fraction of the experimental budget that a factorial design would require. Gradient-boosting tree models have also been trained to predict growth parameters based on medium conditions [Hashizume et al., 2023, Zhang et al., 2023]. Multi-fidelity methods fuse a cheap assay such as growth measurements in microtiter plates with an expensive one, such as cell counting or bioreactor validation [Cosenza et al., 2022, Martens et al., 2025, Sun et al., 2022]. Substituting an *in silico* model trained on experimental data for experimental validation is also proposed without exploring the limitations and evaluation metrics needed for such a substitution [Siska et al., 2026].

Applications studies of this kind demonstrate a loop end to end and verify the winning condition experimentally to evaluate the performance of the loop. However, comparing design strategies is challenging because conditions that the surrogate model inside the closed loop ranks low are never run, which means the counterfactual strategies are not assessed. In practice, the field resolves this by scoring strategies against an *in silico* objective [Siska et al., 2026]. De Luca et al compare iterative model-based strategies across single and parallel reactor set-ups on a fixed simulator, finding that outcomes vary substantially with strategy and that strategies incorporating a model-validity measure outperform those maximising the target quantity alone [De Luca et al., 2023]. Outside bioprocess, benchmarking work in materials science has shown that the surrogate used inside the loop shapes the search path taken, even where different surrogates converge on the same optimum [Liang et al., 2021]. However, a comprehensive evaluation strategy for surrogate verifiers in bioprocess design has not been devised. The impact of the *in silico* surrogate on design strategy selection and model transfer remains unknown.

### 2.2 Verification methods for machine learning systems

AI systems that propose scientific claims can now enumerate and rank candidates at the scale of compute, beyond the limits of manual inspection or experimental testing [Yamada et al., 2025, Mitchener et al., 2025, Gottweis et al., 2026]. As a result, verifying claims made by these systems requires applying statistical techniques that check whether the model’s predictions are reliable enough to act on [Cornelio et al., 2025]. Models in deployment must be qualified within a specified context of use, not by accuracy alone [Coombs et al., 2026]. In this work, we use three quantities from the machine learning literature: the *calibration* of a model’s stated uncertainty, the *coverage* its intervals actually attain, and the *robustness* of both under distribution shift.

Calibration assesses whether a model’s claim about its uncertainty on a prediction holds empirically, by measuring how often the true value falls within its stated interval. For example, a model with nominal 90% intervals that contain the true value 70% of the time is overconfident; one that achieves 99% is conservative. In addition to calibration, model sharpness ranks models by their interval size: narrow intervals are preferable among equally-calibrated models [Gneiting et al., 2007].

Coverage is the probability that a prediction interval contains the true outcome. Conformal prediction provides a model-agnostic guarantee: given a held-out calibration set, it converts any point predictor into intervals attaining a chosen coverage level in finite samples [Fontana et al., 2023, Angelopoulos and Bates, 2023]. Importantly, the guarantee is marginal, which means it holds on average over the population, and is compatible with systematic over-coverage in one region of the input space offset by under-coverage in another. The test and calibration data must be exchangeable and drawn from the same distribution [Foygel Barber et al., 2021].

When models are transferred from one context (such as a microbial strain, experimental scale, or lab setting) to another, the input distribution and input-output relationship can both shift. Iterative design loops introduce a further complication known as feedback covariate shift: because the loop queries areas of the design space where the model predicts favorably, the experimentally validated samples depend on the model itself [Fannjiang et al., 2022]. Coverage guarantees established on one input distribution do not hold on another, which requires further evaluation of the model in the new context. We compute the minimum calibration set required to guarantee coverage on a new domain by evaluating the realized coverage of a transferred model.

## 3 Methods

### 3.1 Data generation and processing

We used three strains, obtained from BCCM: *Lacticaseibacillus rhamnosus* LMG 6400 (Strain A), *Lacticaseibacillus rhamnosus* GG (LMG18243) (Strain B), and *Lacticaseibacillus paracasei* subsp. *paracasei* (LMG12165) (Strain C). We used Latin hypercube sampling to create a DOE of 10 media components (glucose, yeast extract, meat extract, soy peptone, sodium acetate, ammonium citrate, potassium phosphate, manganese sulfate, magnesium sulfate, and Tween 80) on a 5-level grid at the levels given in Table 4. Growth was measured via optical density at 600 nanometers. Runs were incubated at 37°C and automatically read every 30 minutes. A base media was included in all non-control wells to support minimal growth. We inoculated the cultures with 1% inoculum and sealed plates to prevent contamination and facilitate anaerobic growth. Positive MRS rich media controls and PBS negative controls were included on every plate to assess plate-to-plate variability. For a full schematic, see Figure 1.

Following data parsing, raw growth curves were baseline-corrected per well. The baseline per-well is the mean raw optical density (OD) over the first 4 hours corrected to floor at zero (by 0.2). The Gompertz curve, a four-parameter sigmoidal curve commonly used to fit and smooth bacterial growth curve readings,

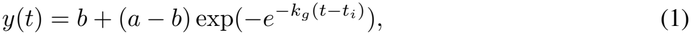

was fitted to each corrected trace by bounded nonlinear least squares [Zwietering et al., 1990]. All reported growth curve metrics are functions of the fitted parameters rather than of individual readings: maximal OD is the upper asymptote *a*, specific growth rate the rate coefficient *k*_*g*_, time of maximal growth the inflection point *t*_*i*_, and lag time is derived from *b, t*_*i*_ and *k*_*g*_. Wells whose fit failed to converge, produced an undefined growth curve parameter, or implied a lag time exceeding the run duration were excluded.

### 3.2 Model training and active learning strategies

We fit six surrogate families to identical wells on all strains: a quadratic response-surface model (ridge regression on second-order polynomial features), a Gaussian process with Matérn-5/2 kernel and per-dimension length scales, random forest, gradient-boosted trees, an ensemble of five multilayer perceptrons, and a mechanistic kinetic model. The mechanistic model was a Monod/Haldane saturation-kinetics model fitted by nonlinear least squares (see Appendix A for details).

The models were chosen to be diverse rather than to maximize accuracy. The kinetic model’s functional form is the only model whose functional form is fixed by mechanism rather than fitted, and the only one that extrapolates substantially above its training wells (1.28× vs. the GP’s 1.09×). Full model hyperparameters are given in Appendix B. Exact Gaussian process inference is cubic in sample size, so to limit fit times the GP is fitted to a random subsample of 2,000 wells.

Wells are partitioned 60/20/20 into training, calibration, and test sets. Following data processing and quality control, there are 2,178 independent compositions shared across all three strains. Partitioning is performed over media compositions, not over wells, so that replicate measurements of one composition are assigned together in the case of strain-to-strain transfer learning. This constraint prevents the same condition from straddling the training and calibration sets and making empirical coverage estimates optimistic. Every reported quantity is the median of 20 independent splits; all seeds derive deterministically from one master seed. Design strategies queried the continuous box spanned by each component’s lowest and highest level; candidates were not snapped to the five experimental levels, so the oracle was also evaluated at concentrations that were never measured. Geometry metrics (roughness, local optima, Sobol indices) were computed on the 5^1^0 experimental lattice.

### 3.3 Design strategy benchmarking

Following surrogate evaluation, each family is then refitted to all Strain C wells and used as the objective - the surrogate oracle - that design strategies are scored against. This follows the offline model-based optimization convention of fitting the evaluation oracle to more data than the task it scores [Trabucco et al., 2022]: here the oracle sees 2,532 wells while each strategy accumulates only 400 oracle queries, so no strategy can reverse-engineer the oracle solely from its own observations.

Design strategies comprise a passive Latin hypercube floor and three Bayesian optimization arms (expected improvement (EI), upper confidence bound (UCB), Thompson sampling (TS)) over a Gaussian process surrogate (see Appendix D for more details). Each cell (oracle, acquisition function, batch size, seed) receives an identical budget of 16 Round-0 wells drawn by Latin hypercube sampling plus 384 adaptive wells, allocated as 384*/q* rounds of *q* wells for *q* ∈1, 2, 4, 8, 16, 32, 64, 96. Within a seed, all arms are dealt the same Round-0 design, so differences between arms are not due to initialization. Note that the optimizers carry their own internal surrogate — the Gaussian process — which is distinct from the oracle they are probing.

Because the families disagree by more than 50% about the attainable optimum, an absolute yield is not comparable across oracles. We therefore report the fraction of each oracle’s own achievable improvement that a strategy captures as the *normalized yield*.

### 3.4 Conformal calibration

We wrap each fitted surrogate in a split conformal predictor [Vovk et al., 2005, Angelopoulos and Bates, 2023]. Nonconformity scores are computed on the held-out calibration partition. At miscoverage level *α*, the intervals are constructed to contain the true value with probability at least 1 *−α*. We evaluate *α* ∈ *α* 0.2, 0.1, 0.05 and report *α* = 0.1 throughout unless stated. Three score functions are compared: the absolute residual; the residual normalised by each model’s native heuristic uncertainty, which yields input-dependent widths; and conformalised quantile regression where the model family supports quantile outputs (excludes quad and Monod families).

Marginal validity is compatible with systematic over-coverage in one region offset by under-coverage in another, so we additionally calibrate group-conditionally (Mondrian conformal), partitioning by predicted-performance decile [Boström et al., 2021]. Decile edges are fixed on the calibration set’s predictions so that a given decile denotes the same region of the response surface on the test set; groups with fewer than 30 calibration points fall back to the global quantile. A calibration set of fewer than 1*/α −*1 points cannot certify *α* at all, in which case the interval is unbounded; the implementation reports such cases as refusals rather than as coverage of one, and they are excluded from all tables.

### 3.5 Transfer protocol

We transferred a LightGBM model trained on the source strain(s) for up to 200 boosting rounds with early stopping. Training is then extended by 100 further rounds on the target strain, freezing the source trees and fitting new trees to the target’s residuals [Ke et al., 2017]. The budget of *k* wells from the target strain is divided evenly between fine-tuning and calibration to model dividing an experimental budget between certifying a model and adapting it. The remaining target-strain wells form the test set; fine-tuning, calibration, and test wells are mutually disjoint and share no compositions. We compare four arms: the source model applied without adaptation (*zero-shot*); the adapted model carrying intervals calibrated on the source strains (*source-cal*); the adapted model recalibrated on target wells (*transfer*); and a model trained from scratch on only the same target wells (*scratch*). We report *k*\*, the smallest k on our grid at which the median realized coverage of the transfer reaches 0.89 and stays above it for every larger *k. k*\* is read off the coverage curve and is not a sample-size guarantee for a future calibration set.

We validated the transfer outside the retrospective split by running a single plate on Strain C. We used a verifier fine-tuned from Strain A on 100 Strain C conditions, frozen before the plate design. Six formulations were selected to span the model’s predicted range (0.63–1.56 OD600) and plated in triplicate; none appeared in any training set. A historical Strain C formulation was included as a bridge control to quantify assay drift.

## 4 Results

### 4.1 Model choice affects prediction of optimal media and design strategy selection

#### Accuracy does not correlate with decision strategy metrics

Across six model families fit on identical data, held-out *R*^2^ spans 0.607 to 0.794. However, accuracy does not correlate with predicted optimum, landscape roughness, or any other decision strategy metric (see Table 1, column 1, 2, 4). Disagreement is not correlated with accuracy; the two most accurate models disagree the most on their decision strategy metrics: GBT (*R*^2^ = 0.794) and RF (*R*^2^ = 0.770) differ by only 3% in *R*^2^, while their believed optima differ by 0.222 and the value they assign to adaptive design differs by 5.3× (0.480 vs. 0.090).

**Table 1:** Accuracy and decision strategy metrics for six models (Strain C, *n* = 2,532). *R*^2^, roughness and local optima are medians over 20 repeats; the optimum and extrapolation ratio come from the oracle refitted to all wells; the batching spread and LHS advantage from the design-strategy sweep. Highest values per column are in **bold**.

| oracle | held-out *R*<sup>2</sup> | optimum (OD) | extrap. vs. max | roughness | local optima | batching spread | adv. over LHS |
|---|---|---|---|---|---|---|---|
| gbt | **0.794** | 2.013 | 0.934 | 0.153 | **122** | **0.546** | 0.480 |
| rf | 0.770 | 1.791 | 0.831 | 0.094 | 30.5 | 0.284 | 0.090 |
| gp | 0.721 | 2.347 | 1.088 | 0.144 | 8.0 | 0.223 | 0.433 |
| nn_ens | 0.719 | 1.865 | 0.865 | 0.161 | 23.0 | 0.200 | **0.586** |
| monod | 0.635 | **2.762** | **1.281** | 0.091 | 21.5 | 0.147 | -0.006 |
| quad | 0.607 | 2.063 | 0.958 | **0.191** | 8.5 | 0.007 | 0.563 |

**Table 2:** Conformal intervals on strain C, *α* = 0.10, absolute nonconformity score, medians over 20 repeats. Coverage is marginal (one global quantile); widths are under Mondrian per-decile calibration, so coverage is equal in every bin and width is the only free quantity.

|  | rf | gbt | nn_ens | gp | monod | quad | spread |
|---|---|---|---|---|---|---|---|
| marginal coverage | 0.905 | 0.903 | 0.906 | 0.905 | 0.906 | 0.908 | 0.005 |
| top-decile width (OD) | **0.251** | 0.306 | 0.363 | 0.419 | 0.465 | **0.778** | 3.1× |
| width range across deciles | 4.3× | 3.1× | 2.9× | 3.0× | 2.7× | 1.9× | — |

**Table 3:**
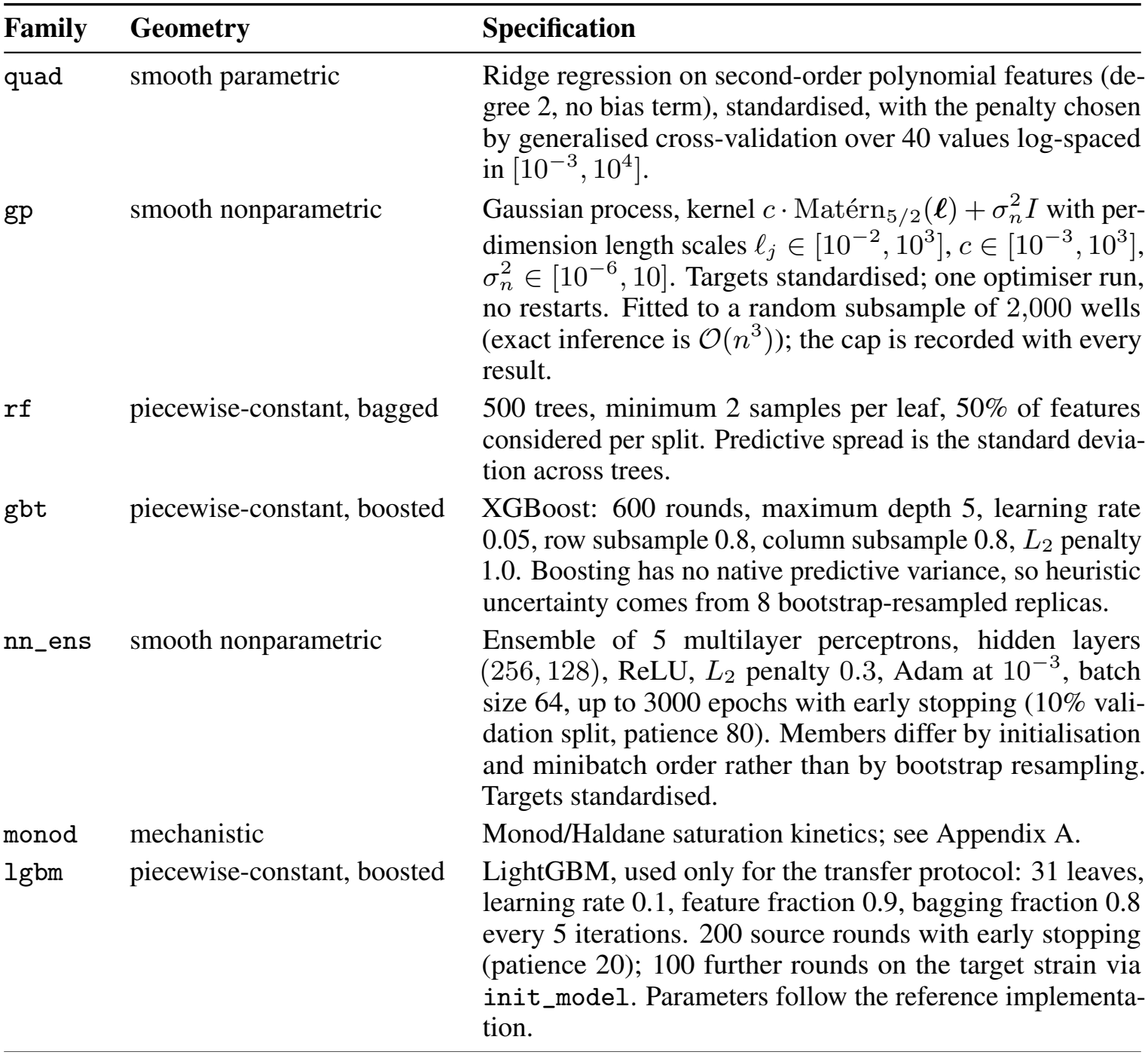
Surrogate families. Families were chosen to span response-surface geometry rather than to maximise accuracy. All values are library defaults except the neural ensemble’s *L*_2_ penalty, selected by a coarse sweep on a single split (see below).

| Family | Geometry | Specification |
|---|---|---|
| quad | smooth parametric | Ridge regression on second-order polynomial features (de-gree 2, no bias term), standardised, with the penalty chosen by generalised cross-validation over 40 values log-spaced in [10<sup>−3</sup>, 10<sup>4</sup>]. |
| gp | smooth nonparametric | Gaussian process, kernel c · Matérn<sub>5/2</sub>(**ℓ**) + σ<sub>n</sub><sup>2</sup>I with per-dimension length scales ℓ<sub>j</sub> ∈ [10<sup>−2</sup>, 10<sup>3</sup>], c ∈ [10<sup>−3</sup>, 10<sup>3</sup>], σ<sub>n</sub><sup>2</sup> ∈ [10<sup>−6</sup>, 10]. Targets standardised; one optimiser run, no restarts. Fitted to a random subsample of 2,000 wells (exact inference is 𝒪(n<sup>3</sup>)); the cap is recorded with every result. |
| rf | piecewise-constant, bagged | 500 trees, minimum 2 samples per leaf, 50% of features considered per split. Predictive spread is the standard devia-tion across trees. |
| gbt | piecewise-constant, boosted | XGBoost: 600 rounds, maximum depth 5, learning rate 0.05, row subsample 0.8, column subsample 0.8, L<sub>2</sub> penalty 1.0. Boosting has no native predictive variance, so heuristic uncertainty comes from 8 bootstrap-resampled replicas. |
| nn_ens | smooth nonparametric | Ensemble of 5 multilayer perceptrons, hidden layers (256, 128), ReLU, L<sub>2</sub> penalty 0.3, Adam at 10<sup>−3</sup>, batch size 64, up to 3000 epochs with early stopping (10% vali-dation split, patience 80). Members differ by initialisation and minibatch order rather than by bootstrap resampling. Targets standardised. |
| monod | mechanistic | Monod/Haldane saturation kinetics; see Appendix A. |
| lgbm | piecewise-constant, boosted | LightGBM, used only for the transfer protocol: 31 leaves, learning rate 0.1, feature fraction 0.9, bagging fraction 0.8 every 5 iterations. 200 source rounds with early stopping (patience 20); 100 further rounds on the target strain via init_model. Parameters follow the reference implementa-tion. |

**Table 4:**
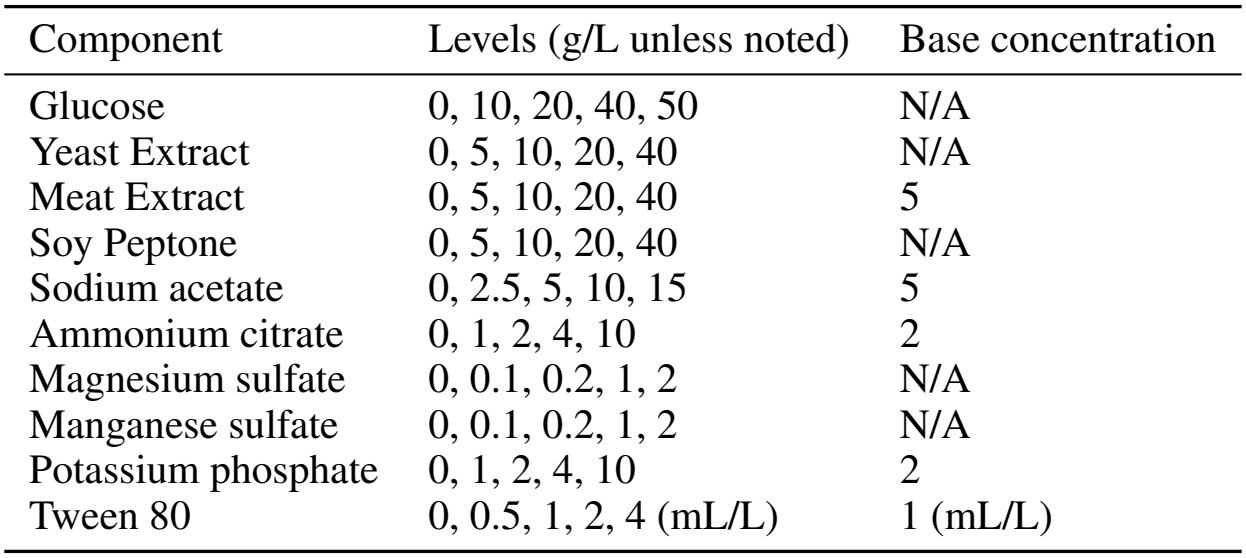
Media components and concentration levels for experimental design and base media.

#### Top components are consistent across models

Total-effect Sobol indices [Tosin et al., 2020] computed on each fitted surface rank glucose first for all six families, and their full ten-component rankings agree at Kendall *τ* = 0.87 − 0.89 on average (minimum 0.73). On Strain C all six return the same top three — glucose, manganese sulfate and yeast extract — in the same order, across polynomial, tree-ensemble and mechanistic forms (Table 7).

#### Models disagree about achievable growth and optimal media

Searched over the design box, the six optima span 1.79–2.76 OD — a 54% range in what is believed attainable from identical data. Relative to the best well ever measured, the random forest’s optimum sits 17% below it while the mechanistic oracle’s sits 28% above. The media they nominate are near-disjoint, with no component appearing in every top-four list.

#### The design decisions a benchmark recommends change with the surrogate

We ran batch Bayesian optimization sweeps against each oracle and assessed how much batch size affects the maximum OD achieved. The batching spread (Table 1, column 6), or difference between the best and worst batch sizes, ranges from 0.7% to over 50%. Batching effects are not monotonic: the GP and mechanistic oracles peak at *q* = 4 and *q* = 2, respectively (see Figure 2a). When comparing to a LHS baseline, adaptive design’s advantage also varies by surrogate: the RF model does not benefit from adaptive sampling, whereas the NN and quad oracles capture 56–59% more of the achievable improvement than LHS. The acquisition function ranking is also affected by the surrogate choice. Thompson sampling performs best on the NN-ensemble and GP oracles, UCB on GBT and RF, and no acquisition function significantly improves on LHS for the Monod oracle (see Figure 2b). This null result is specific to the GP verifier: an RF verifier captures 0.85 of the achievable improvement on Monod (Table 14).

**Figure 2:**
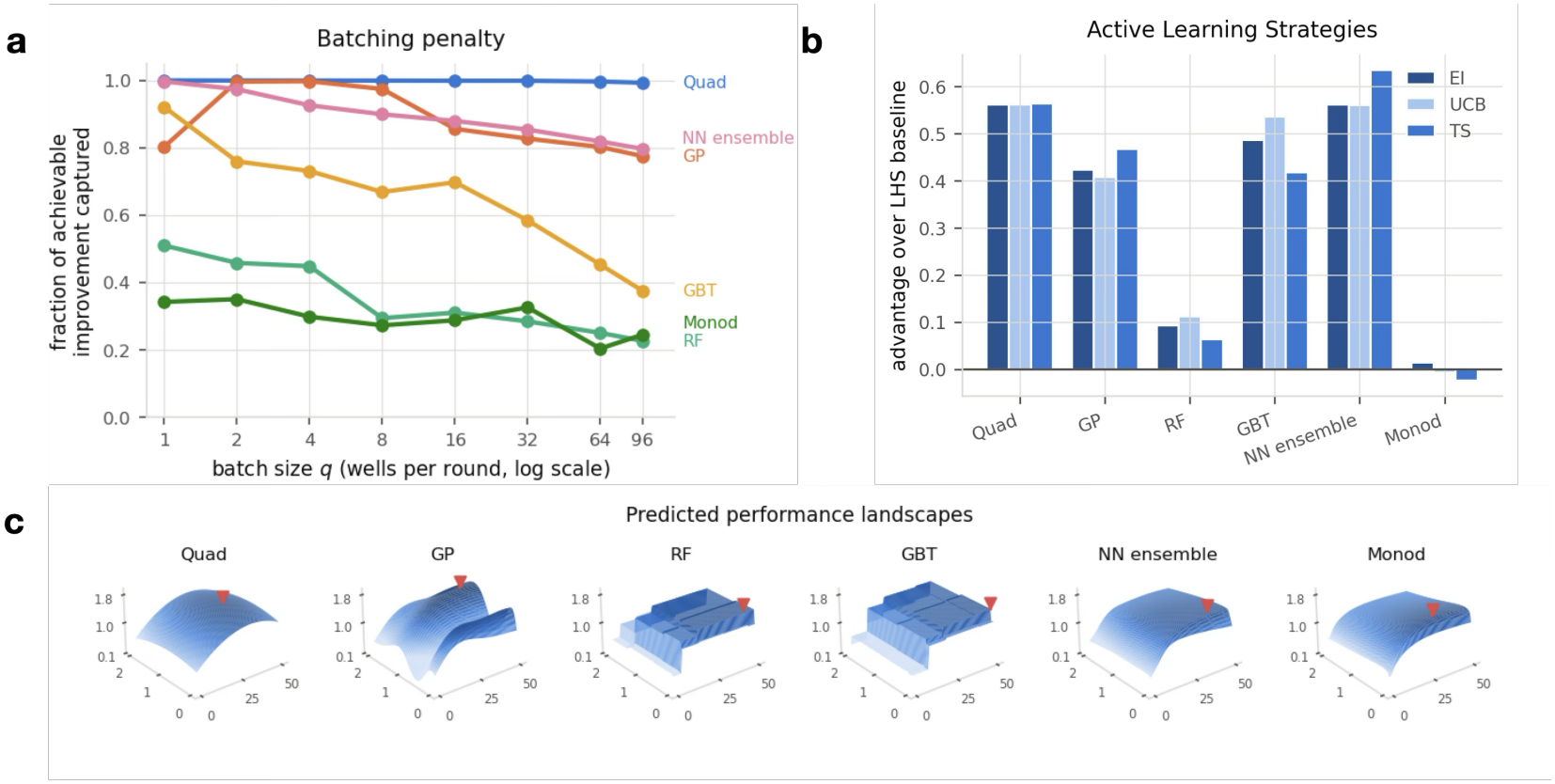
Surrogate oracle family affects closed-loop design strategies. Six model families trained on Strain C (N=2532) to predict maxOD600. **(a)** Batching penalties expressed as a fraction of achievable improvement captured by an expected improvement (EI) Bayesian optimization (BO) loop, stratified by model family. **(b)** Advantage over a Latin-hypercube baseline of 3 acquisition functions (Expected Improvement, *q*-batch Upper Confidence Bound (UCB), and Thompson Sampling (TS) averaged over all batch sizes). **(c)** The surfaces over the two components with the largest total-effect Sobol indices (Glucose and manganese sulfate; other components set to design medians) differ in geometry. Red triangle marks the oracle’s optimum.

#### Design strategy advantage tracks surface geometry

Roughness and advantage over a LHS baseline (see Figure 2c) correlate strongly (Spearman *ρ* = 0.94). Despite being the model with the second-highest accuracy, random forest’s flat surface means that it is accordingly the oracle on which Bayesian optimization barely beats a Latin hypercube (0.090 against 0.586).

### 4.2 Conformal calibration is region-dependent, not model-dependent

#### Interval sharpness correlates with but does not replicate accuracy

At *α*=0.05, 0.10, and 0.20, all six families are statistically indistinguishable on calibration validity (Table 8). All three nonconformity scores behave identically; validity cannot be used to distinguish between models as every family obtains nominal coverage (Table 9). Sharpness separates them: the top-decile widths span 3.1*×*. However, it largely re-states accuracy rather than adding to it (Kendall *τ* = 0.73 between the two rankings), so it does not operate as a separate “trustworthiness” axis for model selection.

**Table 5:**
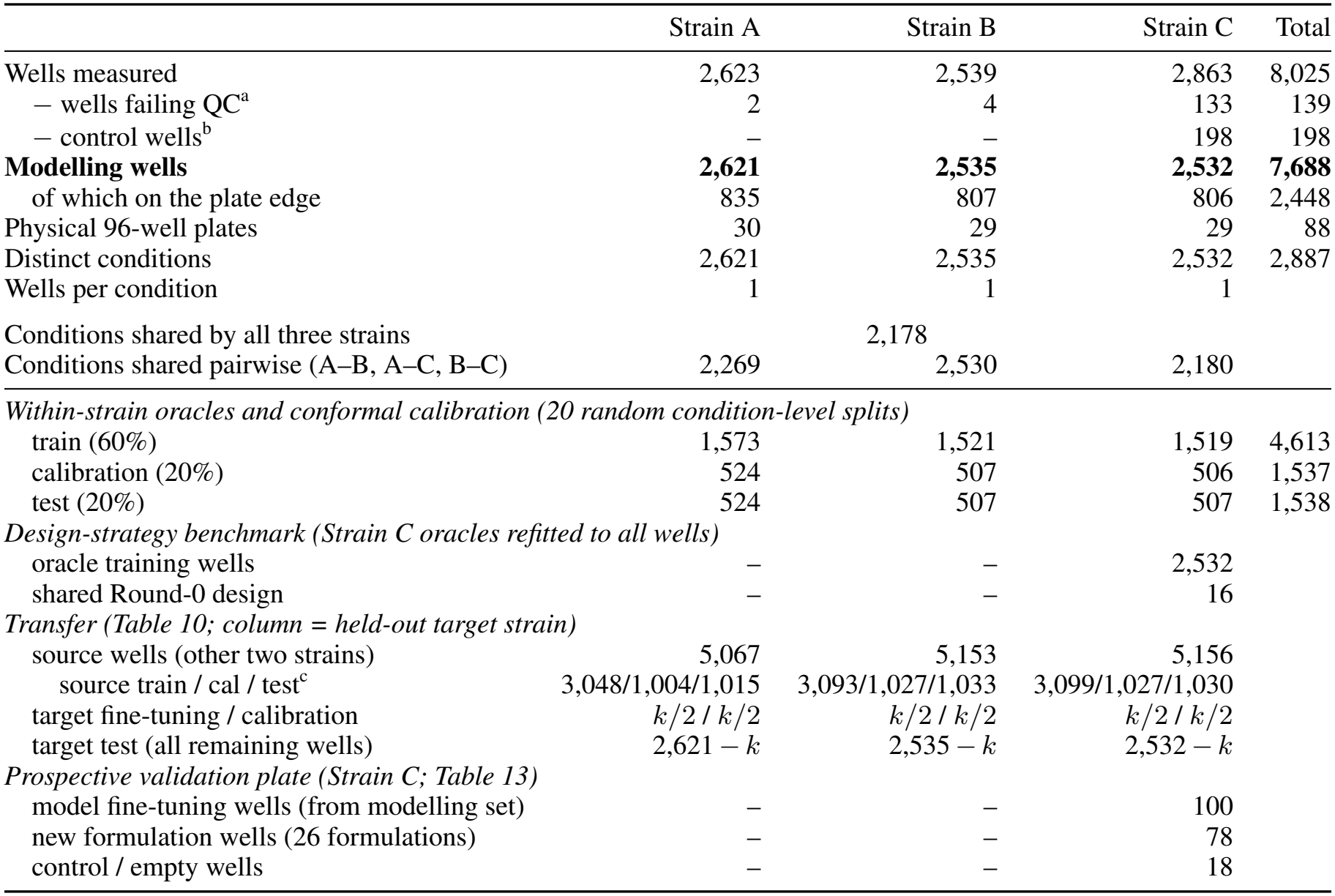
Data accounting, from plate-reader export to the wells each analysis uses. Strain A = *L. rhamnosus* LMG 6400, B = *L. rhamnosus* GG (LMG 18243), C = *L. paracasei* subsp. *paracasei* LMG 12165. Each condition (a distinct 10-component composition) has one well per strain, so all splits are by condition. ^a^Removed by upstream curve-fitting QC. ^b^Column 12 of Strain C’s plates holds medium blanks and controls. ^c^One of the 10 repeats; the others differ by a few wells.

|  | Strain A | Strain B | Strain C | Total |
|---|---|---|---|---|
| Wells measured | 2,623 | 2,539 | 2,863 | 8,025 |
| − wells failing QC<sup>a</sup> | 2 | 4 | 133 | 139 |
| − control wells<sup>b</sup> | – | – | 198 | 198 |
| **Modelling wells** | **2,621** | **2,535** | **2,532** | **7,688** |
| of which on the plate edge | 835 | 807 | 806 | 2,448 |
| Physical 96-well plates | 30 | 29 | 29 | 88 |
| Distinct conditions | 2,621 | 2,535 | 2,532 | 2,887 |
| Wells per condition | 1 | 1 | 1 |  |
| Conditions shared by all three strains | 2,178 |  |  |  |
| Conditions shared pairwise (A–B, A–C, B–C) | 2,269 | 2,530 | 2,180 |  |
| *Within-strain oracles and conformal calibration (20 random condition-level splits)* |  |  |  |  |
| train (60%) | 1,573 | 1,521 | 1,519 | 4,613 |
| calibration (20%) | 524 | 507 | 506 | 1,537 |
| test (20%) | 524 | 507 | 507 | 1,538 |
| *Design-strategy benchmark (Strain C oracles refitted to all wells)* |  |  |  |  |
| oracle training wells | – | – | 2,532 |  |
| shared Round-0 design | – | – | 16 |  |
| *Transfer (Table 10; column = held-out target strain)* |  |  |  |  |
| source wells (other two strains) | 5,067 | 5,153 | 5,156 |  |
| source train / cal / test<sup>c</sup> | 3,048/1,004/1,015 | 3,093/1,027/1,033 | 3,099/1,027/1,030 |  |
| target fine-tuning / calibration | *k*/2 / *k*/2 | *k*/2 / *k*/2 | *k*/2 / *k*/2 |  |
| target test (all remaining wells) | 2,621 − *k* | 2,535 − *k* | 2,532 − *k* |  |
| *Prospective validation plate (Strain C; Table 13)* |  |  |  |  |
| model fine-tuning wells (from modelling set) | – | – | 100 |  |
| new formulation wells (26 formulations) | – | – | 78 |  |
| control / empty wells | – | – | 18 |  |

**Table 6:** Acquisition settings. All arms share one BoTorch `SingleTaskGP` surrogate (RBF-ARD, normalised inputs, standardised outputs, exact MLL).

| arm | implementation | batch construction | settings |
|---|---|---|---|
| lhs | Latin hypercube | independent draw | — |
| gp_ei | qLogExpectedImprovement | joint, optimize_acqf | 10 restarts, 256 raw |
| gp_qucb | qUpperConfidenceBound | joint, optimize_acqf | β = 2, 10 restarts, 256 raw |
| gp_ts | PathwiseThompsonSampling | joint, optimize_acqf | 10 restarts, 256 raw |

**Table 7:** Total-effect Sobol indices computed on each oracle’s own fitted surface, medians over 20 repeats.

| component | gbt | rf | gp | nn_ens | monod | quad |
|---|---|---|---|---|---|---|
| *Strain A (LMG6400)*, *L. rhamnosus* |  |  |  |  |  |  |
| glucose | 0.484 | 0.566 | 0.521 | 0.513 | 0.613 | 0.576 |
| manganese sulfate | 0.275 | 0.273 | 0.335 | 0.211 | 0.191 | 0.169 |
| yeast extract | 0.174 | 0.183 | 0.149 | 0.172 | 0.133 | 0.134 |
| ammonium citrate | 0.156 | 0.117 | 0.176 | 0.153 | 0.014 | 0.137 |
| meat extract | 0.128 | 0.112 | 0.111 | 0.115 | 0.052 | 0.077 |
| soy peptone | 0.026 | 0.009 | 0.010 | 0.034 | 0.007 | 0.022 |
| potassium phosphate | 0.064 | 0.026 | 0.019 | 0.046 | 0.023 | 0.032 |
| magnesium sulfate | 0.049 | 0.029 | 0.042 | 0.039 | 0.003 | 0.036 |
| sodium acetate | 0.024 | 0.005 | 0.001 | 0.019 | 0.004 | 0.019 |
| Tween 80 | 0.012 | 0.003 | 0.000 | 0.012 | 0.002 | 0.005 |
| pairwise Kendall τ: mean 0.89, minimum 0.73 |  |  |  |  |  |  |
| *Strain B (LMG18243)*, *L. rhamnosus* |  |  |  |  |  |  |
| glucose | 0.561 | 0.638 | 0.614 | 0.591 | 0.625 | 0.590 |
| manganese sulfate | 0.150 | 0.153 | 0.178 | 0.096 | 0.099 | 0.080 |
| yeast extract | 0.198 | 0.202 | 0.166 | 0.180 | 0.158 | 0.162 |
| ammonium citrate | 0.061 | 0.041 | 0.052 | 0.052 | 0.014 | 0.049 |
| meat extract | 0.136 | 0.120 | 0.116 | 0.120 | 0.070 | 0.090 |
| soy peptone | 0.049 | 0.019 | 0.040 | 0.052 | 0.035 | 0.050 |
| potassium phosphate | 0.046 | 0.017 | 0.035 | 0.036 | 0.014 | 0.023 |
| magnesium sulfate | 0.040 | 0.018 | 0.017 | 0.033 | 0.005 | 0.030 |
| sodium acetate | 0.031 | 0.007 | 0.015 | 0.029 | 0.010 | 0.029 |
| Tween 80 | 0.009 | 0.002 | 0.000 | 0.012 | 0.002 | 0.005 |
| pairwise Kendall τ: mean 0.87, minimum 0.78 |  |  |  |  |  |  |
| *Strain C (LMG12165)*, *L. paracasei* |  |  |  |  |  |  |
| glucose | 0.646 | 0.741 | 0.710 | 0.690 | 0.754 | 0.703 |
| manganese sulfate | 0.207 | 0.180 | 0.211 | 0.152 | 0.132 | 0.124 |
| yeast extract | 0.120 | 0.111 | 0.095 | 0.103 | 0.083 | 0.094 |
| ammonium citrate | 0.070 | 0.040 | 0.064 | 0.068 | 0.005 | 0.064 |
| meat extract | 0.059 | 0.037 | 0.039 | 0.052 | 0.021 | 0.039 |
| soy peptone | 0.053 | 0.028 | 0.037 | 0.052 | 0.025 | 0.052 |
| potassium phosphate | 0.043 | 0.017 | 0.020 | 0.032 | 0.005 | 0.025 |
| magnesium sulfate | 0.030 | 0.018 | 0.010 | 0.018 | 0.006 | 0.018 |
| sodium acetate | 0.020 | 0.005 | 0.005 | 0.016 | 0.003 | 0.011 |
| Tween 80 | 0.016 | 0.003 | 0.000 | 0.013 | 0.002 | 0.009 |
| pairwise Kendall τ: mean 0.89, minimum 0.73 |  |  |  |  |  |  |

**Table 8:** Marginal coverage of split-conformal intervals, absolute nonconformity score, medians over 20 repeats.

|  | Strain A |  |  | Strain B |  |  | Strain C |  |  |
|---|---|---|---|---|---|---|---|---|---|
| oracle | .95 | .90 | .80 | .95 | .90 | .80 | .95 | .90 | .80 |
| rf | 0.953 | 0.899 | 0.802 | 0.952 | 0.905 | 0.801 | 0.959 | 0.905 | 0.810 |
| gbt | 0.952 | 0.891 | 0.800 | 0.953 | 0.902 | 0.800 | 0.953 | 0.903 | 0.797 |
| nn_ens | 0.956 | 0.901 | 0.805 | 0.959 | 0.908 | 0.812 | 0.963 | 0.906 | 0.800 |
| gp | 0.954 | 0.906 | 0.790 | 0.952 | 0.902 | 0.813 | 0.956 | 0.905 | 0.805 |
| monod | 0.957 | 0.904 | 0.803 | 0.956 | 0.903 | 0.803 | 0.955 | 0.906 | 0.807 |
| quad | 0.957 | 0.906 | 0.798 | 0.955 | 0.908 | 0.809 | 0.957 | 0.908 | 0.808 |

**Table 9:** Marginal coverage at *α* = 0.10 under the three nonconformity scores, Strain C, medians over 20 repeats. The scores are interchangeable for validity. Conformalised quantile regression requires quantile outputs, which the quadratic and mechanistic families cannot produce.

| score | rf | gbt | nn_ens | gp | monod | quad |
|---|---|---|---|---|---|---|
| absolute | 0.905 | 0.903 | 0.906 | 0.905 | 0.906 | 0.908 |
| normalised | 0.904 | 0.913 | 0.905 | 0.904 | 0.897 | 0.904 |
| CQR | 0.903 | 0.907 | 0.901 | 0.904 | — | — |

#### Residual scale varies more across the response surface than between model families

Per-decile calibration targets 90% coverage in each bin, so width differences are not driven by unequal coverage. Interval width varies as much within a family across deciles (1.9–4.3*×*) as between families within a decile (1.2–3.1 *×*; see Table 2). Where a model is asked to predict changes what it can certify more than which model was chosen. Under a single global quantile, this phenomenon results in a systematic error: coverage is only 0.877 in the lowest three deciles and 0.957 in the highest three, where top candidates would be selected for further escalation.

### 4.3 Model calibration does not survive transfer

#### Zero-shot transfer leads to low-coverage

Without fine-tuning on held-out target strain wells, the zero-shot model scores *R*^2^ = 0.713 on held-out source wells and 0.577 on the target strain (see Figure 3a). The conformal intervals, calibrated on source data, drop from 90% coverage to 82.7%.

**Figure 3:**
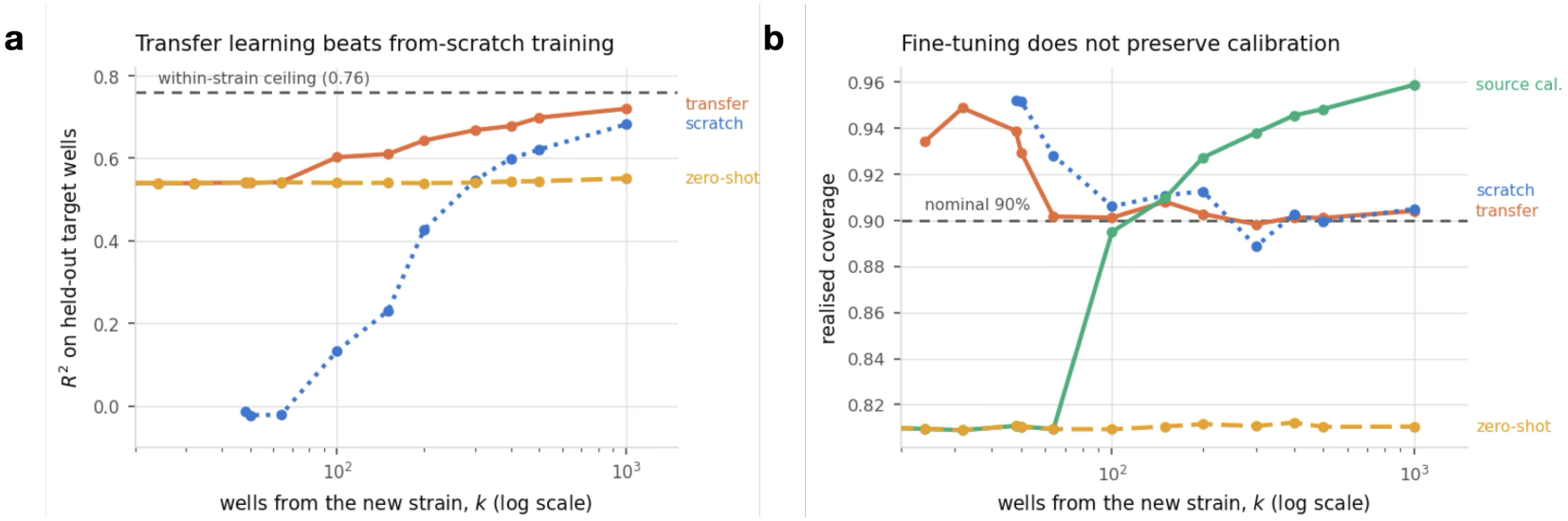
Transfer of models strain-to-strain requires fine-tuning. LightGBMs were trained on leave-one-strain-out source sets and transferred to an unseen target strain (ten seeds per fold, transferred with `init_model` on *k* target wells split evenly between fine-tuning and calibration; median curves over 30 runs). **(a)** Fine-tuning lifts *R*^2^ from 0.57 to 0.73. **(b)** Coverage of 90% calibration intervals; *k <* 18 cannot certify *α* = 0.10 and is omitted. TRANSFER and SOURCE-CAL arms are the same model calibrated on target and held-out source wells, respectively.

However, this marginal figure understates the problem: binned by predicted performance, zero-shot coverage runs from 0.706 in the fourth decile to 0.946 in the ninth.

#### Fine-tuning recovers accuracy; only target well calibration restores coverage

Fine-tuning the model on the target strain recovers accuracy by *k* = 1000 wells; however, calibration does not follow. As fine-tuning moves the model away from the source strain distribution, its residuals there grow. The SOURCE-CAL arm is the *same fitted model* as TRANSFER by construction, differing only in which wells the interval is calibrated on. As the number of calibration wells increases, SOURCE-CAL goes from under-covering intervals below *k* = 100 to over-covering intervals above it (Table 10). In contrast, the TRANSFER arm achieves 90% coverage at *k* = 64 (see Figure 3b).

**Table 10:** Transfer to a held-out strain (LightGBM, medians over 30 runs: 3 held-out strains *×* 10 seeds). TRANSFER and SOURCE-CAL share one fitted model, so *R*^2^ is common to both. *Marginal*: standard split conformal, coverage *≥* 0.90 on average over calibration draws. *PAC*: coverage *≥*0.90 with probability *≥* 0.95 over the calibration draw, from the same calibration scores [Vovk, 2012]; “viol.” is the fraction of runs whose realised coverage fell more than two binomial standard errors of the evaluation set (*≈*0.012) below 0.90. *n*_cal_ = *k/*2 target wells. Widths are measured in OD_600_.

|  |  |  | TRANSFER |  |  |  |  | SOURCE-CAL |  |
|---|---|---|---|---|---|---|---|---|---|
|  |  |  | marginal |  | PAC |  |  | marginal |  |
| *k* | *n*<sub>cal</sub> | *R*<sup>2</sup> | cov. | width | cov. | width | viol. | cov. | width |
| 0 (zero-shot) | — | 0.540 | — | — | — | — | — | 0.810 | 0.706 |
| 64 | 32 | 0.542 | 0.902 | 0.891 | 0.969 | 1.356 | 3% | 0.809 | 0.706 |
| 100 | 50 | 0.603 | 0.901 | 0.863 | 0.960 | 1.164 | 7% | 0.895 | 0.827 |
| 300 | 150 | 0.668 | 0.898 | 0.732 | 0.935 | 0.871 | 7% | 0.938 | 0.907 |
| 1000 | 500 | 0.720 | 0.904 | 0.690 | 0.922 | 0.753 | 0% | 0.959 | 0.958 |

**Table 11:** Optimum, normalisation and design-strategy values behind Table 1. Medians are over 20 seeds per oracle, arm and batch size.

|  | optimum (OD) |  |  |  |  |  | batching (EI) |  | advantage over LHS |  |  |  |
|---|---|---|---|---|---|---|---|---|---|---|---|---|
| oracle | multistart | best BO | ref. | extrap. | Round-0 | achievable | best/worst *q* | spread | EI | UCB | TS | mean |
| gbt | 2.013 | 2.013 | **2.013** | 0.934 | 1.668 | 0.345 | 1/96 | 0.546 | 0.487 | 0.536 | 0.418 | 0.480 |
| rf | 1.791 | 1.762 | **1.791** | 0.831 | 1.628 | 0.163 | 1/96 | 0.284 | 0.093 | 0.112 | 0.064 | 0.090 |
| gp | 2.262 | 2.347 | **2.347** | 1.088 | 1.929 | 0.417 | 4/96 | 0.223 | 0.423 | 0.408 | 0.467 | 0.433 |
| nn_ens | 1.865 | 1.865 | **1.865** | 0.865 | 1.717 | 0.148 | 1/96 | 0.200 | 0.562 | 0.560 | 0.635 | 0.586 |
| monod | 2.762 | 2.674 | **2.762** | 1.281 | 1.740 | 1.022 | 2/64 | 0.147 | 0.014 | -0.008 | -0.024 | -0.006 |
| quad | 2.063 | 2.065 | **2.065** | 0.958 | 1.848 | 0.216 | 1/96 | 0.007 | 0.562 | 0.562 | 0.564 | 0.563 |

**Table 12:**
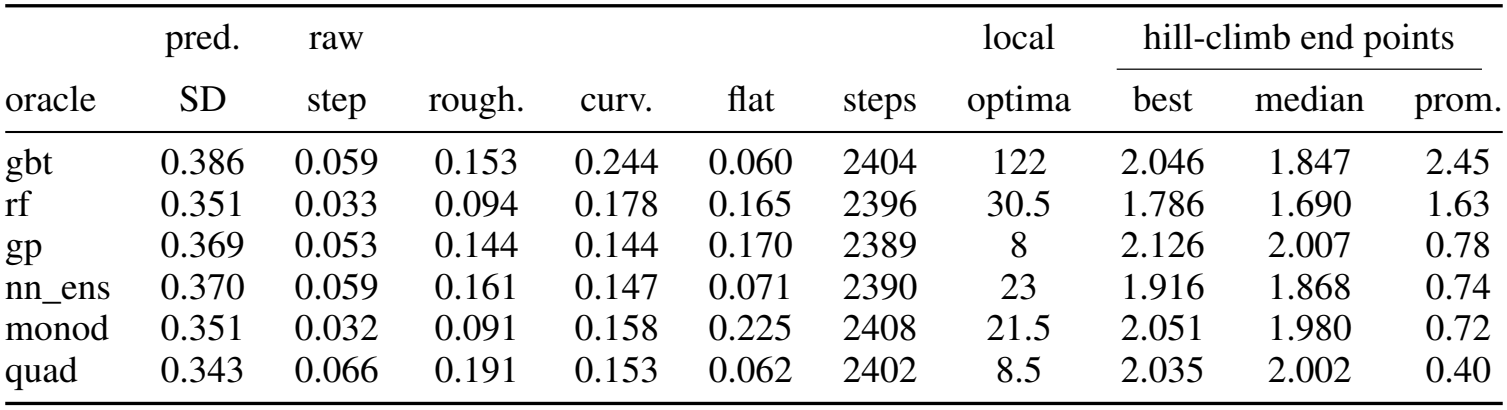
Surface geometry for models in Table 1. For definitions of metrics, see E.1

**Table 13:**
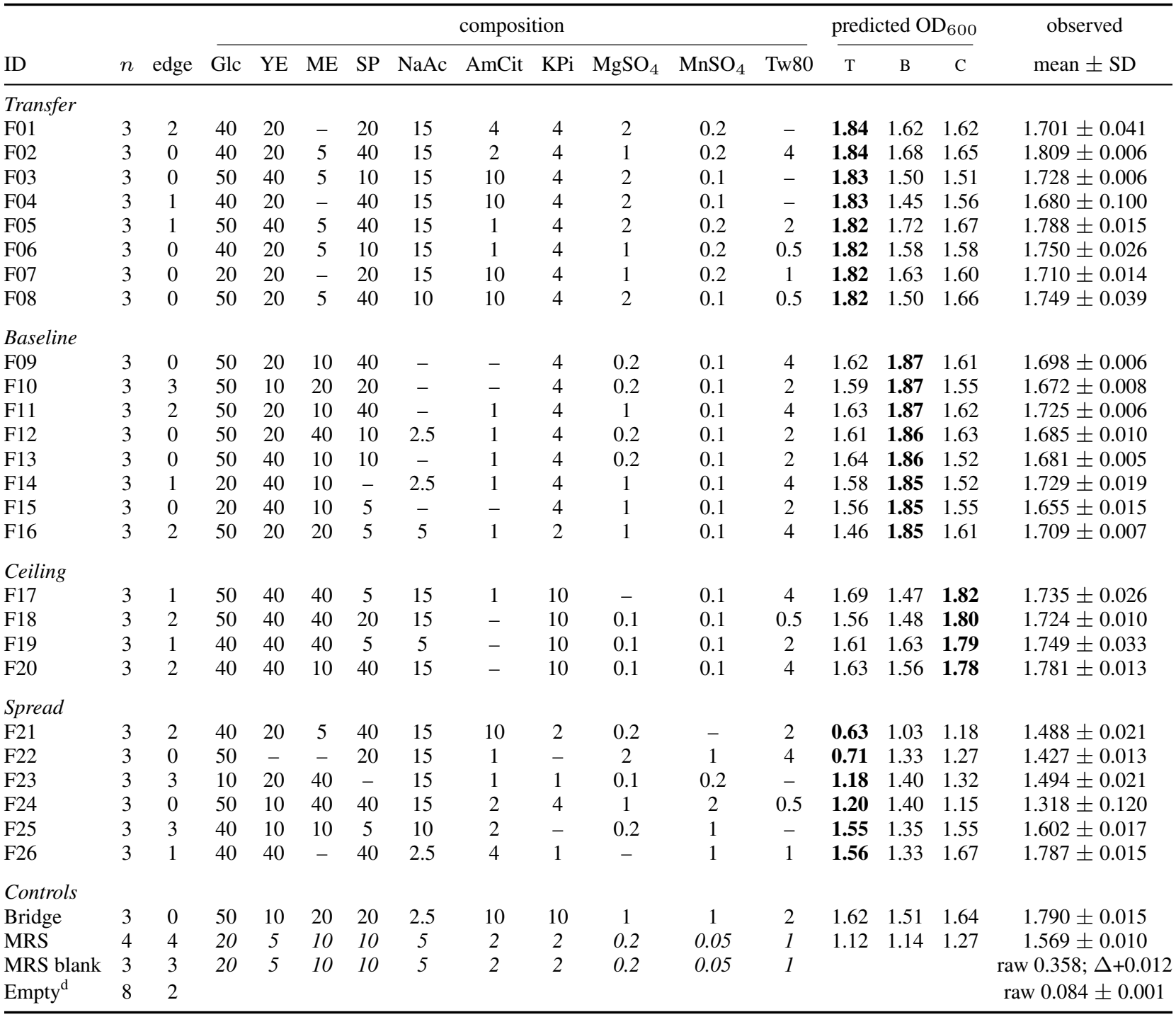
Prospective validation plate for LightGBM model transfer on Strain C. *Baseline* and Transfer formulations are the top-predicted formulations for the 100-well model and its fine-tuned correlate. *Spread* formulations were chosen by T to span its predicted range.

|  |  |  | composition |  |  |  |  |  |  |  |  |  | predicted OD<sub>600</sub> |  |  | observed |
|---|---|---|---|---|---|---|---|---|---|---|---|---|---|---|---|---|
| ID | *n* | edge | Glc | YE | ME | SP | NaAc | AmCit | KPi | MgSO<sub>4</sub> | MnSO<sub>4</sub> | Tw80 | T | B | C | mean ± SD |
| *Transfer* |  |  |  |  |  |  |  |  |  |  |  |  |  |  |  |  |
| F01 | 3 | 2 | 40 | 20 | – | 20 | 15 | 4 | 4 | 2 | 0.2 | – | **1.84** | 1.62 | 1.62 | 1.701 ± 0.041 |
| F02 | 3 | 0 | 40 | 20 | 5 | 40 | 15 | 2 | 4 | 1 | 0.2 | 4 | **1.84** | 1.68 | 1.65 | 1.809 ± 0.006 |
| F03 | 3 | 0 | 50 | 40 | 5 | 10 | 15 | 10 | 4 | 2 | 0.1 | – | **1.83** | 1.50 | 1.51 | 1.728 ± 0.006 |
| F04 | 3 | 1 | 40 | 20 | – | 40 | 15 | 10 | 4 | 2 | 0.1 | – | **1.83** | 1.45 | 1.56 | 1.680 ± 0.100 |
| F05 | 3 | 1 | 50 | 40 | 5 | 40 | 15 | 1 | 4 | 2 | 0.2 | 2 | **1.82** | 1.72 | 1.67 | 1.788 ± 0.015 |
| F06 | 3 | 0 | 40 | 20 | 5 | 10 | 15 | 1 | 4 | 1 | 0.2 | 0.5 | **1.82** | 1.58 | 1.58 | 1.750 ± 0.026 |
| F07 | 3 | 0 | 20 | 20 | – | 20 | 15 | 10 | 4 | 1 | 0.2 | 1 | **1.82** | 1.63 | 1.60 | 1.710 ± 0.014 |
| F08 | 3 | 0 | 50 | 20 | 5 | 40 | 10 | 10 | 4 | 2 | 0.1 | 0.5 | **1.82** | 1.50 | 1.66 | 1.749 ± 0.039 |
| *Baseline* |  |  |  |  |  |  |  |  |  |  |  |  |  |  |  |  |
| F09 | 3 | 0 | 50 | 20 | 10 | 40 | – | – | 4 | 0.2 | 0.1 | 4 | 1.62 | **1.87** | 1.61 | 1.698 ± 0.006 |
| F10 | 3 | 3 | 50 | 10 | 20 | 20 | – | – | 4 | 0.2 | 0.1 | 2 | 1.59 | **1.87** | 1.55 | 1.672 ± 0.008 |
| F11 | 3 | 2 | 50 | 20 | 10 | 40 | – | 1 | 4 | 1 | 0.1 | 4 | 1.63 | **1.87** | 1.62 | 1.725 ± 0.006 |
| F12 | 3 | 0 | 50 | 20 | 40 | 10 | 2.5 | 1 | 4 | 0.2 | 0.1 | 2 | 1.61 | **1.86** | 1.63 | 1.685 ± 0.010 |
| F13 | 3 | 0 | 50 | 40 | 10 | 10 | – | 1 | 4 | 0.2 | 0.1 | 2 | 1.64 | **1.86** | 1.52 | 1.681 ± 0.005 |
| F14 | 3 | 1 | 20 | 40 | 10 | – | 2.5 | 1 | 4 | 1 | 0.1 | 4 | 1.58 | **1.85** | 1.52 | 1.729 ± 0.019 |
| F15 | 3 | 0 | 20 | 40 | 10 | 5 | – | – | 4 | 1 | 0.1 | 2 | 1.56 | **1.85** | 1.55 | 1.655 ± 0.015 |
| F16 | 3 | 2 | 50 | 20 | 20 | 5 | 5 | 1 | 2 | 1 | 0.1 | 4 | 1.46 | **1.85** | 1.61 | 1.709 ± 0.007 |
| *Ceiling* |  |  |  |  |  |  |  |  |  |  |  |  |  |  |  |  |
| F17 | 3 | 1 | 50 | 40 | 40 | 5 | 15 | 1 | 10 | – | 0.1 | 4 | 1.69 | 1.47 | **1.82** | 1.735 ± 0.026 |
| F18 | 3 | 2 | 50 | 40 | 40 | 20 | 15 | – | 10 | 0.1 | 0.1 | 0.5 | 1.56 | 1.48 | **1.80** | 1.724 ± 0.010 |
| F19 | 3 | 1 | 40 | 40 | 40 | 5 | 5 | – | 10 | 0.1 | 0.1 | 2 | 1.61 | 1.63 | **1.79** | 1.749 ± 0.033 |
| F20 | 3 | 2 | 40 | 40 | 10 | 40 | 15 | – | 10 | 0.1 | 0.1 | 4 | 1.63 | 1.56 | **1.78** | 1.781 ± 0.013 |
| *Spread* |  |  |  |  |  |  |  |  |  |  |  |  |  |  |  |  |
| F21 | 3 | 2 | 40 | 20 | 5 | 40 | 15 | 10 | 2 | 0.2 | – | 2 | **0.63** | 1.03 | 1.18 | 1.488 ± 0.021 |
| F22 | 3 | 0 | 50 | – | – | 20 | 15 | 1 | – | 2 | 1 | 4 | **0.71** | 1.33 | 1.27 | 1.427 ± 0.013 |
| F23 | 3 | 3 | 10 | 20 | 40 | – | 15 | 1 | 1 | 0.1 | 0.2 | – | **1.18** | 1.40 | 1.32 | 1.494 ± 0.021 |
| F24 | 3 | 0 | 50 | 10 | 40 | 40 | 15 | 2 | 4 | 1 | 2 | 0.5 | **1.20** | 1.40 | 1.15 | 1.318 ± 0.120 |
| F25 | 3 | 3 | 40 | 10 | 10 | 5 | 10 | 2 | – | 0.2 | 1 | – | **1.55** | 1.35 | 1.55 | 1.602 ± 0.017 |
| F26 | 3 | 1 | 40 | 40 | – | 40 | 2.5 | 4 | 1 | – | 1 | 1 | **1.56** | 1.33 | 1.67 | 1.787 ± 0.015 |
| *Controls* |  |  |  |  |  |  |  |  |  |  |  |  |  |  |  |  |
| Bridge | 3 | 0 | 50 | 10 | 20 | 20 | 2.5 | 10 | 10 | 1 | 1 | 2 | 1.62 | 1.51 | 1.64 | 1.790 ± 0.015 |
| MRS | 4 | 4 | *20* | *5* | *10* | *10* | *5* | *2* | *2* | *0.2* | *0.05* | *1* | 1.12 | 1.14 | 1.27 | 1.569 ± 0.010 |
| MRS blank | 3 | 3 | *20* | *5* | *10* | *10* | *5* | *2* | *2* | *0.2* | *0.05* | *1* |  |  |  | raw 0.358; Δ+0.012 |
| Empty<sup>d</sup> | 8 | 2 |  |  |  |  |  |  |  |  |  |  |  |  |  | raw 0.084 ± 0.001 |

**Table 14:** Verifier *×*oracle cross (Strain C, *q* = 8, 384 adaptive wells, 10 seeds). Median normalised yield, as in Table 1. Every verifier row uses the same acquisition (analytic EI on a 2,048-point Sobol pool with local-search polish). Family-matching effect (*δ* in yield = verifier + oracle + *δ·* matched) +0.108 [+0.076, +0.142] (95% seed bootstrap).

|  | oracle |  |  |  |  |  |
|---|---|---|---|---|---|---|
| verifier | gbt | rf | gp | nn_ens | monod | quad |
| GP | 0.541 | 0.395 | 0.782 | 0.845 | 0.585 | 0.960 |
| RF | 0.450 | 0.513 | 0.635 | 0.592 | 0.847 | 0.665 |
| MLP ens. | 0.549 | 0.417 | 0.707 | 0.815 | 0.677 | 0.936 |
| GP-EI (joint) | 0.618 | 0.271 | 0.979 | 0.894 | 0.345 | 0.999 |
| LHS | 0.157 | 0.242 | 0.431 | 0.352 | 0.291 | 0.436 |

#### Certifying a new organism requires several 96-well plates of experiments

Defining *k*^*⋆*^ as the smallest budget at which median coverage reaches 0.89 and does not drop below it again, *k*^*⋆*^ = 100, 400 and 300 wells for the three held-out strains. In addition to the variation between strains, *k*^*⋆*^ certifies only the marginal claim. The coverage spread across performance deciles drops from 0.240 at zero-shot to 0.090 at *k* = 48, but does not consistently converge to a low value; at *k* = 100, spread rises to 0.174 (Table 10).

#### Validating certification failures prospectively

We tested the transferred verifier on six formulations spread across the predicted range. The model’s 90% intervals contained four of the six formulations, and both misses were the two lowest-predicted formulations: media predicted at 0.63 and 0.71 OD600 grew to 1.49 and 1.43. Underprediction scaled with how poor the model believed a medium to be. Six formulations cannot establish a coverage rate, but the direction and size of the failures match the retrospective decile analysis.

## 5 Conclusion

Closed-loop experimental design requires a clear understanding of a model’s accuracy, uncertainty, and failure modes. As the number of AI-driven experimental candidates increases, verification of model decisions grows in importance. Scarce wet-lab confirmation budgets limit the assessability of many different model arms; in this work, we instead investigate the model surrogate in an active learning loop, using conformal verification, and under conditions of distribution shift and model transfer. We find that global accuracy is an underdetermined metric for model selection; decision strategies, extrapolation ability, and landscape geometry all vary per-surrogate family, and the media they nominate are near-disjoint. We recommend running decisions across several oracles to ensure candidate robustness. We also find that transfer to a new unseen strain requires an experimental budget for both fine-tuning (to recover model accuracy) and calibration (to ensure coverage). Verifying AI-driven predictions is not solely a question of whether it is accurate enough, but of whether the conclusion drawn from it would survive a different, equally defensible model. Conformal calibration, extrapolation ratio, and landscape geometry metrics are all cheap to measure, and should be reported alongside accuracy when a surrogate is qualified for a stated context of use.

### Limitations

All models were trained on three strains from the same genus, across the same limited medium condition design space, for one prediction target (growth). We anticipate that the magnitude of surrogate-to-surrogate variability will vary across new species and experimental assays. This work supports the choice of surrogate as an underexamined variable in closed-loop design, but does not establish which oracle or design strategy gives the best outcome across real experimental cases. Additionally, the transfer-calibration analysis is performed using only one model, LightGBM, and additional experimental validation of surrogate-specific candidates would solidify the paper’s claims. The required experimental budget *k*^*⋆*^ varies across strain, likely due to experimental noise and metabolic differences, but with only three strains, we cannot say how that budget scales to a larger range of less-related organisms. Furthermore, we do not assess the effect of changing the split between fine-tuning and calibration (fixed at 0.5 throughout); *k*^*⋆*^ could be an artifact of that allocation.

### Recommendations

The machine learning surrogate in an active learning loop should be considered a design variable and tested as such. A surrogate’s inductive biases should match not only the underlying problem but also the objective of the acquisition function. Strategies that target global accuracy require assumptions that hold across the entire design space. Strategies that aim to locate the optimum quickly (e.g. EI) only need to model the path towards it accurately, and global accuracy does not reveal whether a surrogate is suited for this. Smooth, conservative surfaces can even hinder the search, as observed for random forests. Unsurprisingly, verifiers in an active learning loop fit oracles from their same mathematical family better (see Table 14), so being aware of the interplay between the surrogate oracle in the loop and the verifier model is important.

### Future work

Several extensions follow directly. First, integrating a model into an iterative design loop induces a covariate shift; the distribution of training inputs from each subsequent iteration are a function of the model used to design the experiments. We recommend applying localized and weighted conformal methods to address this covariate shift. Second, the wet-lab experiments we take as ground truth have noise and batch-to-batch effects. We assessed plate and edge effects, corrected each plate to its negative control baseline, and ran all experiments in per-strain batches to reduce this, but future work could use replicates and more complex control design to more effectively assess noise. Third, disagreement across a surrogate zoo is cheap to compute and could be an acquisition signal rather than just a diagnostic, indicating that regions where oracles disagree the most are areas where a new experiment would most reduce uncertainty.

## Author contributions

CM trained all models in the paper, performed the conformal analysis and developed the transfer learning pipeline. JG designed the Bayesian optimization loop. JC and MP designed the experimental setup and supervised all wet-lab experiments. CM created figures and drafted paper text; JG helped edit paper in response to reviews. The following team members also contributed to running the wet-lab experiments: Kaitlyn Przydzial, Luis Kramer, Kristiana Ellen, Nihan Kardan.

## A Mechanistic Monod-kinetics model

### A.1 Model specification

Let 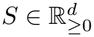be a medium composition, *d* = 10, scaled by each component’s observed maximum,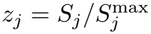. Each component enters through a Haldane term, floored for numerical stability:

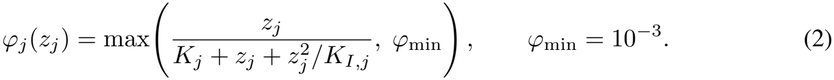

As *K*_*I,j*_ *→ ∞* this reduces to the Monod form *z*_*j*_ */*(*K*_*j*_ + *z*_*j*_); for finite *K*_*I,j*_ it is non-monotone, peaking at 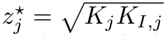 with 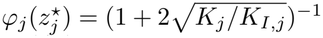, so a component can be limiting at low concentration and inhibitory at high.

Components combine as a weighted geometric mean with an exponential interaction term, evaluated in log space:

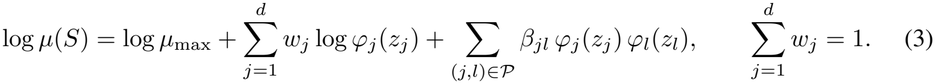

The unit-sum constraint is load-bearing. An unweighted productΠ _*j*_ *φ*_*j*_ decays geometrically in *d*, so with ten components the response would collapse toward zero regardless of composition; the weighted geometric mean keeps *µ* bounded independently of dimension. The floor *φ*_min_ is likewise a modelling choice and not only a numerical guard: a geometric mean vanishes if any factor is zero, and the design deliberately includes a zero level for every component, so without the floor any medium omitting a component would predict no growth. The floor caps the penalty for an omitted component at *w*_*j*_ log *φ*_min_, a soft substitutability assumption.

### A.2 Parameterization and estimation

Parameters are fitted in an unconstrained space so that positivity and the simplex constraint hold by construction,

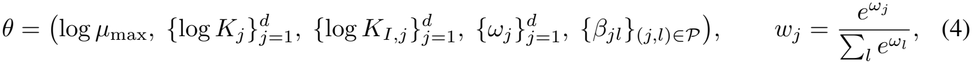

and estimated by nonlinear least squares, 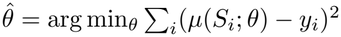, solved by trust-region reflective optimisation (at most 4000 function evaluations) from three random initialisations, retaining the lowest-cost solution. Initialisation draws *K*_*j*_ ~ *U* (0.05, 0.5) and *K*_*I j*_ *~ U* (1, 20) in scaled units, *ω*_*j*_ *~ N* (0, 0.1^2^), *β*_*jl*_ = 0, and 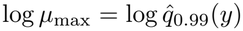. Numerical guards clip log *K*_*j*_ to [−12, 8],log *K*_*I,j*_ to [−12, 12] and log *µ* to [−30, 30]. Half-saturation constants are reported in original units as 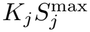.

### A.3 Interaction terms

*P*contains |*P*|= 10 of the 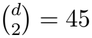 unordered pairs, drawn uniformly without replacement and fixed by seed, giving 41 free parameters. Because the pairs are sampled rather than selected, we checked whether the draw is load-bearing. Across five pair-set seeds, held-out *R*^2^ is stable (0.628–0.644) and the extrapolation ratio exceeds one in every draw.

## B Surrogate model specifications and hyperparameters

All families are fitted to the same wells under the same splits. Inputs are *z*-scored before fitting except where noted; the mechanistic model is fitted in concentration units scaled by each component’s observed maximum, and the LightGBM transfer model on raw concentrations, since tree splits are scale-invariant and raw units keep the fitted model comparable with the reference implementation.

### Hyperparameter selection

Only one hyperparameter was tuned. The neural ensemble’s *L*_2_ penalty was chosen by a coarse sweep over 10^−3^, {10^−2^, 10^−1^, 0.3, 1, 3} on one split of the strain B data; the default of 10^−3^ costs approximately 0.13 held-out *R*^2^ here, and an under-tuned family would weaken the comparison the paper rests on. All other families use library defaults.

## C Experimental conditions

## D Acquisition functions

All three adaptive arms share one surrogate, so any difference between them is the acquisition rule alone. The surrogate is a BoTorch `SingleTaskGP` (v0.17.2) with inputs normalised to the unit cube and outputs standardised, an RBF kernel with one length scale per component (ARD) under BoTorch’s dimension-scaled priors, fitted by exact marginal likelihood. Write *µ*(*x*) and *σ*(*x*) for its posterior mean and standard deviation, *f* ^*⋆*^ for the best value observed so far, and *q* for the batch size.

### Expected improvement (gp_ei)

Score a point by how much it is expected to beat the incumbent, counting only improvements:

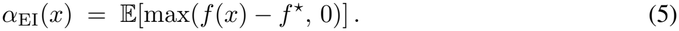

We use the *q*-batch logarithmic form, `qLogExpectedImprovement` [Ament et al., 2023], which optimises log *α* to avoid the vanishing gradients that make plain EI hard to optimise once most candidates are unpromising. Batches are chosen jointly by `optimize_acqf` (10 restarts, 256 raw samples).

### Upper confidence bound (gp_qucb)

Score a point by an optimistic estimate of its value, trading mean against uncertainty through a single parameter:

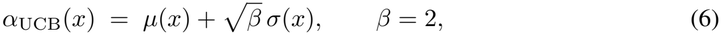

so a point is judged on its posterior mean plus 1.41 standard deviations. The batch version is the Monte-Carlo reparameterisation of Wilson et al. [2017], `qUpperConfidenceBound`, optimised jointly with the same settings as EI. *β* is held fixed rather than annealed, so the explore–exploit balance does not drift over rounds.

### Thompson sampling (gp_ts)

Rather than score points, draw *q* independent functions from the GP posterior and take the maximiser of each:

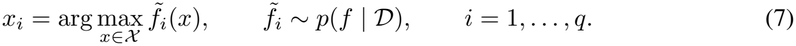

Each 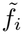is a whole posterior sample path, drawn by Matheron’s rule [Wilson et al., 2020]: a prior function, approximated with random Fourier features of the RBF-ARD kernel, plus an exact data-dependent update that conditions it on the observations. Because a path is a deterministic function once drawn, it can be evaluated anywhere in the continuous design box *X* and maximised with gradients. The *q* paths are drawn once per round and maximised jointly by `optimize_acqf` with the same budget as EI and UCB (10 restarts, 256 raw samples). The *i*-th batch point only enters the *i*-th path, so the joint optimisation maximises each draw separately. Diversity within the batch comes from the sampling itself: different draws place their maxima in different regions in proportion to the posterior probability that the optimum lies there, so no explicit diversity penalty is needed.

### Space-filling floor (lhs)

Latin hypercube sampling over the design box, drawn once per round, ignoring all observations. It is the reference against which the adaptive arms are scored: an arm that cannot beat it has not earned the cost of fitting a model.

## E Supplementary Results Tables

### E.1 Definitions of surface geometry metrics

All metrics are evaluated on the 5^10^ experimental lattice for Strain C and reported as medians over 20 split fits.

- *Pred. SD*: standard deviation of the oracle’s predictions over the lattice; the normalising scale for roughness, curvature and flatness.
- *Raw step*: mean |*f* (*x* + Δ_*j*_) *− f* (*x*) |in OD for a one-level step Δ_*j*_ in one randomly chosen component *j*, at 4,000 random lattice points (interior levels only).
- *Roughness*: raw step divided by pred. SD.
- *Curvature*: mean |*f* (*x* +Δ_*j*_) − 2*f* (*x*) + *f* (*x −* Δ_*j*_)| divided by pred. SD, at the same points.
- *Flat*: fraction of one-level steps that change the prediction by less than 1% of pred. SD.
- *Steps*: number of one-level steps evaluated for the metrics above.
- *Local optima*: number of distinct end points reached by 200 best-improvement hill climbs over lattice neighbours (*±*1 level in one component, *≤*60 steps each).
- *Best*: highest predicted value (OD) among the hill-climb end points.
- *Median*: median predicted value (OD) of the hill-climb end points.
- *Prominence*: (best − median)*/*SD of the end-point values.

## Notes

### Competing Interest Statement

All authors are employees of Differential Bio GmBH and performed this research as part of their employment.

## References

Sebastian Ament, Samuel Daulton, David Eriksson, Maximilian Balandat, and Eytan Bakshy. Unexpected improvements to expected improvement for bayesian optimization. Advances in neural information processing systems, 36:20577–20612, 2023.

Anastasios N Angelopoulos and Stephen Bates. Conformal prediction: A gentle introduction. Foundations and Trends in Machine Learning, 16(4):494–591, 2023.

Henrik Boström, Ulf Johansson, and Tuwe Löfström. Mondrian conformal predictive distributions. In Conformal and Probabilistic Prediction and Applications, pages 24–38. PMLR, 2021.

Wenduo Cheng, Mingqian Ma, Shuaike Shen, Anna Hupalowska, Jennifer E Rood, Yang Zhang, Gaurav Agrawal, Christine Bakan, Michelle A Lee, Aviv Regev, et al. Towards human-led, agent-driven autonomous laboratories for the life sciences. 2026.

Evan Claes, Tommy Heck, Kathleen Coddens, Maarten Sonnaert, Jan Schrooten, and Jan Verwaeren. Bayesian cell therapy process optimization. Biotechnology and Bioengineering, 121(5):1569–1582, 2024.

Dana J Coombs, Bryan Fritz, Ellie Galliker, and Juri Steiner. Application of the asme v&v 40 standard to computational models. Computational Modeling and Simulation of Medical Devices: Verification, Validation, and Uncertainty Quantification, page 325, 2026.

Cristina Cornelio, Takuya Ito, Ryan Cory-Wright, Sanjeeb Dash, and Lior Horesh. The need for verification in ai-driven scientific discovery. arXiv preprint arXiv:2509.01398, 2025.

Zachary Cosenza, Raul Astudillo, Peter I Frazier, Keith Baar, and David E Block. Multi-information source bayesian optimization of culture media for cellular agriculture. Biotechnology and bioengineering, 119(9):2447–2458, 2022.

Riccardo De Luca, Goncalo Costa, Harini Narayanan, Claus Wirnsperger, Mariano N Cruz Bournazou, Alessandro Butte, and Moritz Von Stosch. Comparison of strategies for iterative model-based upstream bioprocess development with single and parallel reactor set-ups. Biochemical Engineering Journal, 191:108813, 2023.

Clara Fannjiang, Stephen Bates, Anastasios N Angelopoulos, Jennifer Listgarten, and Michael I Jordan. Conformal prediction under feedback covariate shift for biomolecular design. Proceedings of the National Academy of Sciences, 119(43):e2204569119, 2022.

Matteo Fontana, Gianluca Zeni, and Simone Vantini. Conformal prediction: a unified review of theory and new challenges. Bernoulli, 29(1):1–23, 2023.

Rina Foygel Barber, Emmanuel J Candes, Aaditya Ramdas, and Ryan J Tibshirani. The limits of distribution-free conditional predictive inference. Information and Inference: A Journal of the IMA, 10(2):455–482, 2021.

Tilmann Gneiting, Fadoua Balabdaoui, and Adrian E Raftery. Probabilistic forecasts, calibration and sharpness. Journal of the Royal Statistical Society Series B: Statistical Methodology, 69(2): 243–268, 2007.

Juraj Gottweis, Wei-Hung Weng, Alexander Daryin, Tao Tu, Petar Sirkovic, Artiom Myaskovsky, Grzegorz Glowaty, Felix Weissenberger, Alessio Orlandi, Dan Popovici, et al. Accelerating scientific discovery with co-scientist. Nature, pages 1–3, 2026.

Takamasa Hashizume, Yuki Ozawa, and Bei-Wen Ying. Employing active learning in the optimization of culture medium for mammalian cells. npj systems biology and applications, 9(1):20, 2023.

Laura Marie Helleckes, Sebastian Putz, Kshitiz Gupta, Matthias Franzreb, and Hector Garcia Martin. Perspectives for artificial intelligence in bioprocess automation. Current Opinion in Biotechnology, 97:103392, 2026.

Masih Karimi Alavijeh, Yih Yean Lee, and Sally L Gras. A perspective-driven and technical evaluation of machine learning in bioreactor scale-up: A case-study for potential model developments. Engineering in Life Sciences, 24(7):e2400023, 2024.

Guolin Ke, Qi Meng, Thomas Finley, Taifeng Wang, Wei Chen, Weidong Ma, Qiwei Ye, and Tie-Yan Liu. Lightgbm: A highly efficient gradient boosting decision tree. Advances in neural information processing systems, 30, 2017.

Shohei Kitano, Ciai Lin, Jee Loon Foo, and Matthew Wook Chang. Synthetic biology: learning the way toward high-precision biological design. PLoS Biology, 21(4):e3002116, 2023.

Frédéric M Lapierre, Pasquale Mattaliano, Dennis Raith, Mariela Castillo-Cota, Jonas Bermeitinger, and Robert Huber. Multi-cycle high-throughput growth media optimization using batch bayesian optimization. Journal of Chemical Technology & Biotechnology, 100(8):1571–1583, 2025.

Qiaohao Liang, Aldair E Gongora, Zekun Ren, Armi Tiihonen, Zhe Liu, Shijing Sun, James R Deneault, Daniil Bash, Flore Mekki-Berrada, Saif A Khan, et al. Benchmarking the performance of bayesian optimization across multiple experimental materials science domains. npj Computational Materials, 7(1):188, 2021.

Adrian Martens, Mathias Neufang, Alessandro Butté, Moritz von Stosch, Antonio del Rio Chanona, and Laura Marie Helleckes. Holistic bioprocess development across scales using multi-fidelity batch bayesian optimization. arXiv preprint arXiv:2508.10970, 2025.

Ludovico Mitchener, Angela Yiu, Benjamin Chang, Mathieu Bourdenx, Tyler Nadolski, Arvis Sulovari, Eric C Landsness, Daniel L Barabasi, Siddharth Narayanan, Nicky Evans, et al. Kosmos: An ai scientist for autonomous discovery. arXiv preprint arXiv:2511.02824, 2025.

Harini Narayanan, Joshua A Hinckley, Rachel Barry, Brendan Dang, Lenna A Wolffe, Adel Atari, Yuen-Yi Tseng, and J Christopher Love. Accelerating cell culture media development using bayesian optimization-based iterative experimental design. Nature communications, 16(1):6055, 2025.

Tien Dung Pham, Robert Bassett, and Uwe Aickelin. Capturing uncertainty in black-box chromatography modelling using conformal prediction and gaussian processes. Computers & Chemical Engineering, 199:109136, 2025.

Maximilian Siska, Emma Pajak, Katrin Rosenthal, Antonio del Rio Chanona, Eric von Lieres, and Laura M Helleckes. A guide to bayesian optimization in bioprocess engineering. Biotechnology and bioengineering, 123(4):805–830, 2026.

Yuan Sun, Winton Nathan-Roberts, Tien Dung Pham, Ellen Otte, and Uwe Aickelin. Multifidelity gaussian process for biomanufacturing process modeling with small data. arXiv preprint arXiv:2211.14493, 2022.

Michel Tosin, Adriano MA Côrtes, and Americo Cunha. A tutorial on sobol’global sensitivity analysis applied to biological models. Networks in systems biology: applications for disease modeling, pages 93–118, 2020.

Brandon Trabucco, Xinyang Geng, Aviral Kumar, and Sergey Levine. Design-bench: Benchmarks for data-driven offline model-based optimization. In International Conference on Machine Learning, pages 21658–21676. PMLR, 2022.

Vladimir Vovk. Conditional validity of inductive conformal predictors. In Asian conference on machine learning, pages 475–490. PMLR, 2012.

Vladimir Vovk, Alexander Gammerman, and Glenn Shafer. Algorithmic learning in a random world. Springer, 2005.

James Wilson, Viacheslav Borovitskiy, Alexander Terenin, Peter Mostowsky, and Marc Deisenroth. Efficiently sampling functions from gaussian process posteriors. In International conference on machine learning, pages 10292–10302. PMLR, 2020.

James T Wilson, Riccardo Moriconi, Frank Hutter, and Marc Peter Deisenroth. The reparameterization trick for acquisition functions. arXiv preprint arXiv:1712.00424, 2017.

Yutaro Yamada, Robert Tjarko Lange, Cong Lu, Shengran Hu, Chris Lu, Jakob Foerster, Jeff Clune, and David Ha. The ai scientist-v2: Workshop-level automated scientific discovery via agentic tree search. arXiv preprint arXiv:2504.08066, 2025.

Shuyang Zhang, Honoka Aida, and Bei-Wen Ying. Employing active learning in medium optimization for selective bacterial growth. Applied Microbiology, 3(4):1355–1369, 2023.

Marcel H Zwietering, Ida Jongenburger, Frank M Rombouts, and KJAEM Van’t Riet. Modeling of the bacterial growth curve. Applied and environmental microbiology, 56(6):1875–1881, 1990.

